# Increases in vein length compensate for leaf area lost to lobing in grapevine

**DOI:** 10.1101/2022.03.15.484490

**Authors:** Zoë Migicovsky, Joel F. Swift, Zachary Helget, Laura L. Klein, Anh Ly, Matthew Maimaitiyiming, Karoline Woodhouse, Anne Fennell, Misha Kwasniewski, Allison J. Miller, Peter Cousins, Daniel H. Chitwood

## Abstract

**Premise:** There is considerable variation in leaf lobing and leaf size, including among grapevines, some of the most well-studied leaves. We examined the relationship between leaf lobing and leaf size across grapevine populations which varied in extent of leaf lobing.

**Methods:** We used homologous landmarking techniques to measure 2,632 leaves across two years in 476 unique, genetically distinct grapevines from 5 biparental crosses which vary primarily in the extent of lobing. We determined to what extent leaf area could explain variation in lobing, vein length, and vein to blade ratio.

**Results:** Although lobing was the primary source of variation in shape across the leaves we measured, leaf area varied only slightly as a function of lobing. Rather, leaf area increases as a function of total major vein length, total branching vein length, and decreases as a function of vein to blade ratio. These relationships are stronger for more highly lobed leaves, with the residuals for each model differing as a function of distal lobing.

**Conclusions:** For a given leaf area, more highly lobed leaves have longer veins and higher vein to blade ratios, allowing them to maintain similar leaf areas despite increased lobing. These findings show how more highly lobed leaves may compensate for what would otherwise result in a reduced leaf area, allowing for increased photosynthetic capacity through similar leaf size.

## Introduction

Each leaf on a plant is shaped by genetics, the environment, and development, which all interact to contribute to variation in form and size (Chitwood and Sinha, 2016). Across diverse species, paleobotanists have long-recognized that more entire (less lobed) leaves are associated with warmer, wetter climates and more toothed leaves, those with serrations around the leaf margins, are found in colder, drier climates (Bailey and Sinnott, 1916; Wilf, 1997). These relationships have been confirmed in extant populations (Peppe et al., 2011; Schmerler et al., 2012; Wright et al., 2017).

Why this variation in lobing and teeth exists across different climates remains an ongoing area of study, with many possible explanations available (Nicotra et al., 2011; Edwards et al., 2016). For example, toothed species examined in a colder region increased transpiration and photosynthate production early in the growing season in comparison to the untoothed species. As a result, this mechanism could provide an advantage to the toothed species by allowing them to maximize the duration of their growing season (Royer and Wilf, 2006). Teeth may also serve as hydathodes, expelling water in early spring in temperate species and avoiding mesophyll flooding due to root pressure (Feild et al., 2005). Additional explanations include that temperate leaves are thinner and rely on structural support from veins, forming a wedge shape and leading to a toothy margin (Givnish, 1979), or that highly dissected leaves reduce feeding efficiency by herbivores (Brown and Lawton, 1991). The shape of the temperate leaf may also be due to selection on leaf primordia inside overwintering buds, with the bounded space resulting in physical pressures influencing the adult leaf form (Edwards et al., 2016). Indeed, similar shapes can arise for many reasons, and the explanations behind lobing across climates may be a combination of these reasons and others.

In addition to leaf lobing, there is considerable variation in leaf size, with an over 100,000-fold difference in leaf size among species worldwide (Díaz et al., 2016; Wright et al., 2017). Similar to variation in lobing, leaf size varies with climate, with larger leaves generally found in wetter, warmer areas, the same zones where more entire leaves are found (Webb, 1968; Peppe et al., 2011; Chitwood and Sinha, 2016). For example, in a study of the Australian rainforest, leaf size was found to decrease with decreasing rainfall (Webb, 1968). However, variation in leaf size has tradeoffs: larger leaves have a thicker boundary layer of still air which slows heat loss and may increase respiration rates more than photosynthesis rates. Additional water is required to cool the leaf by transpiration. At the same time, larger leaves have more photosynthetic potential due to their larger surface area. Thus, when access to water decreases, smaller leaves are favored (Givnish, 1987; Westoby et al., 2002).

Recent efforts to estimate leaf area found that in addition to leaf length and width, accurate estimates required a leaf shape specific correction factor. The correction factor differed depending on the extent of lobing of the leaf, indicating that leaves differed allometrically– differences in size were correlated with other differences in shape, in this case, lobing. More highly lobed leaves had reduced leaf areas in comparison to unlobed leaves with the same dimensions (Schrader et al., 2021).

Indeed, many plant leaves show allometric relationships, which occur as a result of lobing as well as other aspects of leaf shape. For example, across a diverse collection of 869 apples (*Malus* spp.) differences in the length-to-width ratio were the primary source of variation, with variation in the width of the blade, not the length, being significantly correlated with variation in leaf shape (Migicovsky et al., 2018). Allometric relationships for leaf length-to-width ratio have also been described in numerous other plant species, including *Passiflora* (Chitwood and Otoni, 2017), *Solanum* (Chitwood et al., 2013), and *Vitis* (Klein et al., 2017). A recent study of *Vitis* species measured across four years and multiple nodes of development identified vein to blade ratio as an allometric variation of leaf size. As leaf size increased, the proportion of the leaf area composed of blade exponentially increased, while the proportion composed of vein area decreased (Chitwood et al., 2021).

Among the most well-studied leaves are those belonging to grapevine species (*Vitis*), with an entire field of ampelography (“vine” + “writing”) dedicated to their study. Ampelography was first described by Louis Ravaz (1902), and brought to widespread attention for its use in wine grapes by Pierre Galet (1979). The use of ampelography has continued in contemporary times through use of comprehensive morphometric techniques, in particular homologous landmarks (Chitwood et al., 2014; Chitwood, Klein, et al., 2016; Klein et al., 2017; Chitwood, 2021; Harris et al., 2021). Homologous landmarking is based on the shared morphology across grapevine leaves including the presences of seven major veins. Despite numerous studies characterizing the shape of grapevine leaves across species (Chitwood, Klein, et al., 2016), development (Chitwood, Klein, et al., 2016), and years (Chitwood, Rundell, et al., 2016; Chitwood et al., 2021), questions remain. In grapevine, lobing is a major source of variation in leaf shape (Galet, 1979; Chitwood, 2021). Lobing permits sunlight to permeate the canopy of the grapevine, which provides desirable benefits to grape growers, but if more highly lobed leaves are smaller in leaf area this would also reduce photosynthetic capacity of the vine.

Here, we measure a total of 2,632 leaves across two years in 476 unique, genetically distinct grapevine accessions from 5 biparental crosses which vary primarily in the extent of lobing. We demonstrate that leaves of varying sizes do not differ in lobing, but rather, more highly lobed leaves of the same area show increases in leaf vein length and vein to blade ratio. These findings provide evidence for a mechanism in which more lobed leaves maintain similar leaf surface area compared to more entire leaves, despite lobing.

## Materials and Methods

### Sampling

Leaves were sampled from seedlings of five biparental *Vitis* populations located in Madera County, California. 500 seedlings were planted in the vineyard. 450 seedlings shared a seed parent, DVIT 2876. The remaining 50 seedlings had DVIT 2876 as a grandparent. DVIT 2876 ‘Olmo b55-19’ is a compound-leafed accession from the USDA-ARS National Clonal Germplasm repository, suspected to include *Vitis piasezkii* Maximowicz, as one of its parents (or grandparents). The populations were created to examine variation in leaf lobing. The vines were composed of 125 individuals from a DVIT 2876 x unnamed *Vitis vinifera* selection cross (Pop1), 100 individuals from a DVIT 2876 x a different unnamed *Vitis vinifera* selection cross (Pop2), 150 individual from a DVIT 2876 x unnamed *Vitis* hybrid cross (Pop3), 75 individual from a DVIT 2876 x a different unnamed *Vitis* hybrid cross (Pop4), and 50 individuals from a seedling (DVIT 2876 x unnamed *Vitis vinifera* selection) x DVIT 3374 (*Vitis mustangensis* Buckley) cross (Pop5).

The vines sampled were planted in 2017. They were trained to a unilateral cordon and spur pruned. Leaf samples were collected on June 22 and July 12 2018, then again in 2019 on June 14, 19, and July 4. Across the sampling dates within a given year, a total of three mature, representative leaves were sampled from each of the vines and placed into labeled plastic bags. The plastic bags were stored in a cooler during collection and scanned, vein side down, later the same day using a flatbed scanner. Files were saved as JPEGs and named using the accession ID. All original scans used in this study are available from *dryad submission in progress*.

### Landmarking

Leaves were analyzed using 21 landmarks as previously described by (Chitwood, Rundell, et al., 2016; Bryson et al., 2020; Chitwood et al., 2021). These landmarks were placed by manually using ImageJ on one side of the leaf (Abràmoff et al., 2004). The resulting table was saved as a text file with the coordinates for all landmarks and then visualized using ggplot2 v.3.3.5 (Wickham, 2016) in R v.4.1.0 (R Core Team, 2021) to detect mistakes. If the resulting visualization did not look like a leaf, indicating that the landmarking had been performed incorrectly, the landmarking was performed again. The resulting data excluded leaves that were damaged and could not be landmarked, as well as vines that were too small to sample from.

### Data analysis

The resulting text files from each scan were merged in R. In addition, we determined the image size for each scan using the imager package version 0.42.10 in R (Barthelme, 2021) and used ImageJ (Abràmoff et al., 2004) to calculate the number of pixels per cm for all subsequent area calculations. Total leaf area as well as blade and vein areas were calculated using the shoelace algorithm, which calculates the area of a polygon using the landmarks as vertices, following previously described methods (Chitwood et al., 2021). In addition, we calculated the ratio of vein to blade area.

The degree of distal and proximal lobing of each leaf was calculated by first calculating the length of the distal and proximal veins. To perform this calculation, we used the midpoint of the landmarks at the base of the vein and the landmark at the tip of the lobe to calculate the distance between the two points. Similarly, we calculated the distance between the distal and proximal sinus and the landmark at the base of the leaf. Distal lobing values were calculated as the ratio of the length of the distal sinus to the length of the distal lobe, with values increasing as lobing decreases and the same ratio was calculated for proximal lobing. In addition, we calculated the length of the remaining major vein (the midvein) as well as the three branching veins. Total major vein length was calculated as the length of the midvein + the length of the distal vein + the length of the proximal vein. Similarly, total branching vein length was the sum of all three branching veins (Figure 1B).

**Figure 1.**
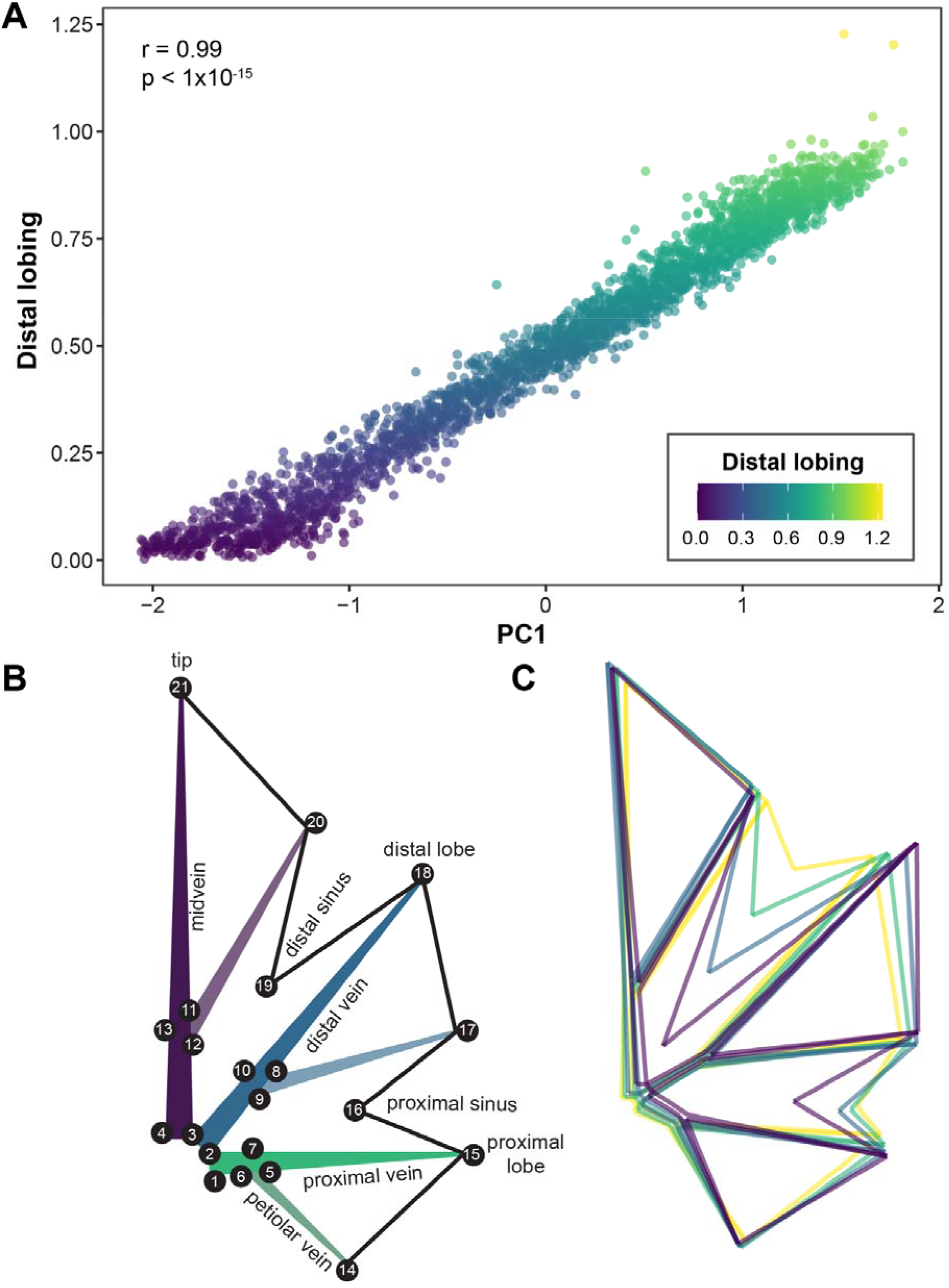
(A) Correlation between principal component 1 (PC1) and distal lobing. Each leaf (n=2632) is plotted with the color of the point indicating the distal lobing value. Distal lobing values are calculated as the ratio of the length of the distal sinus to the length of the distal lobe terminus, with values increasing as lobing decreases. The Pearson’s correlation coefficient between PC1 and distal lobing is indicated, as is the amount of variance explained by PC1. (B) Twenty one landmarks used in this study for measuring leaf shape and size. Major veins are labeled, with branching veins indicated by lighter shades. Numbers indicate the order the landmarks were placed. (C) For each PC1 quartile, a mean leaf is plotted and coloured according to distal lobing value.

Landmarks were adjusted using a Generalized Procrustes Analysis (GPA) in the shapes R package version 1.2.6 with the reflect=TRUE option (Dryden, 2021) before performing principal components analysis (PCA). To compare differences in size across years, we used a Mann-Whitney U test to contrast 2018 and 2019 leaf areas for vines which were fully sampled (3 leaves) in both years..

Subsequent analyses were performed in R and code analyzing the data is available at the following github repository https://github.com/zoemigicovsky/grape_leaf_lobing. Visualizations were performed using ggplot2 (Wickham, 2016). Briefly, for accessions with three leaves sampled in both 2018 and 2019, we examined variation in leaf area, blade area, and vein area as well as the ratio between vein area and blade area for each leaf across both years. We used a Pearson’s correlation to determine the relationship between distal lobing and principal component 1 (PC1) based on homologous landmark data, as well as examining the variation in distal lobing for each of the five crosses sampled in this study.

We modeled the natural logarithm of area (cm^2^) vs distal lobing as well as the natural logarithm of area (cm^2^) vs natural logarithm of total major vein length (cm). We extracted the residuals from the model describing the relationship between ln(area)~ln(total major vein length) and modeled the relationship between the residuals vs distal lobing. Next, we calculated the correlation between total major vein length and total branching vein length, before modeling the relationship between the natural logarithm of area (cm^2^) and natural logarithm of total branching vein length (cm) as well as the residual values vs distal lobing. Lastly, we repeated the analysis examining the relationship between the natural logarithm of area (cm^2^) vs the natural logarithm of the ratio of vein area to blade area, and then modeling the relationship between the residuals from that model and distal lobing.

## Results

For each genetically distinct accession a total of 1-6 leaves were sampled across the two years of the study. In order to compare the shape and size of leaves across the two years, for this analysis only, only accessions with three leaves sampled for both years were included. The total number of accessions with three leaves sampled in both years was 398, for a total of 2388 leaves. Across these 398 accessions, area was significantly greater for the whole leaf (W = 584071, p = 2.14 × 10^−14^) as well as blade (W = 585202, p = 3.59 × 10^−14^) and vein (W = 571944, p < 1 × 10^−15^) regions in 2019 compared to 2018 (Figure S1A). While the median leaf area for 2018 was 24.5 cm^2^, it increased by 4.2 cm^2^ resulting in a median leaf area of 28.7 cm^2^ in 2019. Since proportionally more of the leaf is composed of blade than vein, the increase in blade area contributes more to the increase in overall leaf area than the increase in vein area does. The blade area increased by a median value of 4.1 cm^2^, while the median increase in vein area was only 0.23 cm^2^. The ratio of vein to blade ratio did not differ significantly between years (W = 720044, p = 0.67), although the larger leaves in 2019 had a slightly lower vein to blade ratio (0.0603) in comparison to 2018 (0.0607) (Figure S1B). Thus, differences in leaf size across years were not correlated with differences in the vein to blade ratio, indicating that leaf expansion occurred proportionally for both blade and veins.

To determine the primary source of variation in leaf shape, all 2,632 leaves sampled across 476 vines in 2018 and 2019 were used in all downstream analysis. First, we performed PCA using the homologous landmark data. The primary axis of variation, PC1, explained approximately 43.7% of the variation in the dataset. PC1 was also significantly correlated with distal lobing (r = 0.99, p <1 ×10^−15^; Figure 1). In contrast, proximal lobing was significantly correlated with PC1, but correlation was weaker (r = 0.66, p <1 ×10^−15^). From this analysis, it is evident that distal lobing is the largest source of shape variation. There was also substantial variation in distal lobing across accessions from each of the five populations (Figure S2). For all subsequent analyses, the complete set of 2,632 leaves was also used.

After calculating the total area of each leaf we modeled the natural log of leaf area as a function of distal lobing (Figure 2). While the natural log of total leaf area varied significantly as a function of distal lobing (p = 0.026), the adjusted R^2^ was 0.003, indicating that the effect of distal lobing on total leaf area is small, and the shape of the leaf is decoupled from its overall area.

**Figure 2.**
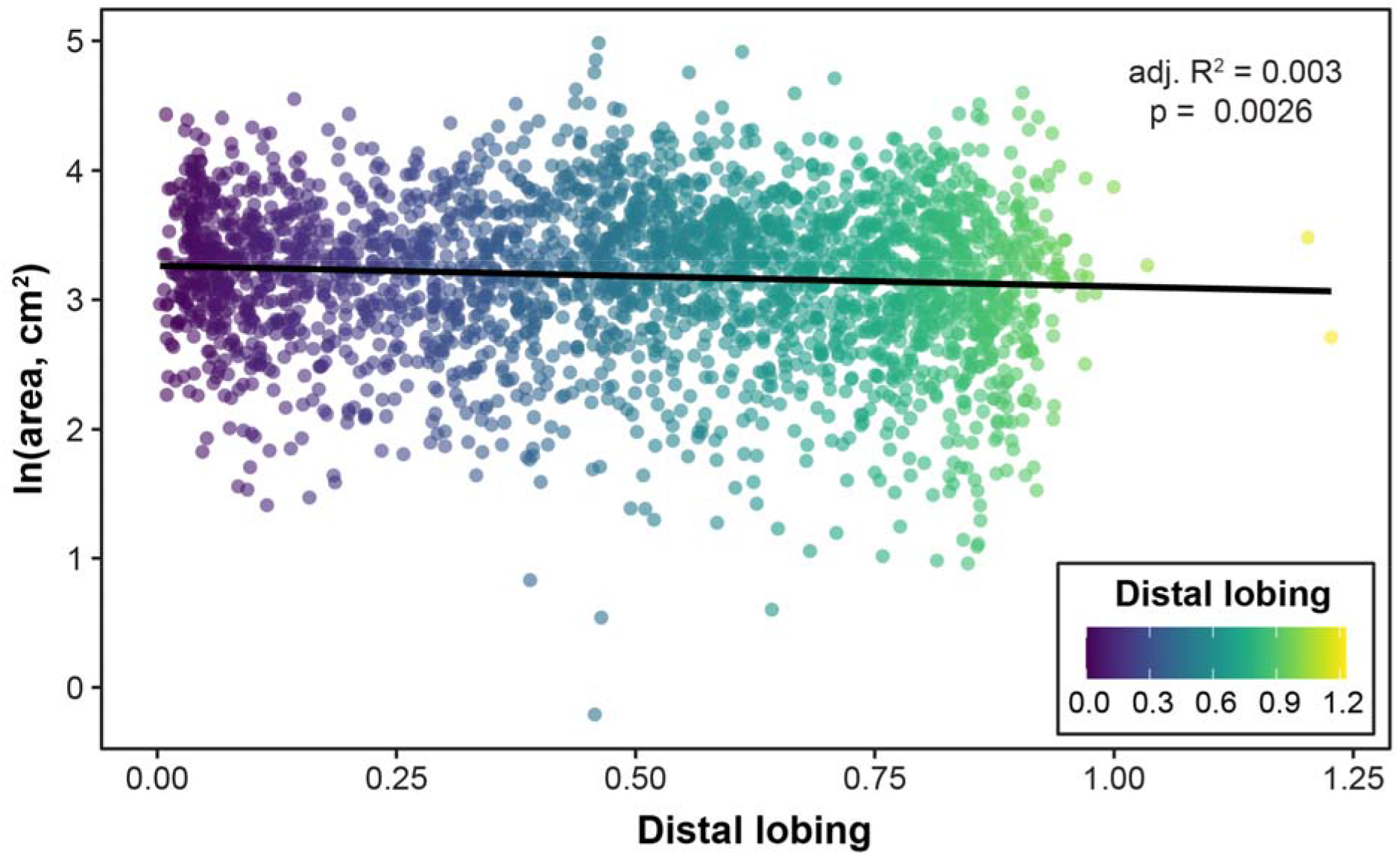
Natural logarithm of area (cm^2^) vs distal lobing. Each leaf (n=2632) is plotted with the color of the point indicating the distal lobing value. The natural log of leaf area varies significantly (p = 0.026) as a function of distal lobing, with an adjusted R^2^ of 0.003.

Modeling the natural log of leaf area as a function of the natural log of total major vein length (the sum of the midvein, distal vein, and proximal vein lengths, indicated in Figure 1B), total leaf area increases as major vein length increases (Figure 3A). This is a linear relationship (R^2^=0.95, p <1 ×10^−15^) indicating that leaf area increases as a function of leaf length. When examining the residuals from this linear model, they show a linear relationship with distal lobing (Figure 3B, R^2^=0.32, p <1 ×10^−15^). Thus, a highly lobed leaf of the same area as a more entire leaf will have longer veins. By increasing vein length and leaf dimensions, highly lobed leaves can maintain similar total leaf areas as more entire leaves. Indeed, by visualizing the mean leaf for each PC1 quartile (Figure 1C), it is apparent that the distal lobe is longer in the more highly lobed leaves.

**Figure 3.**
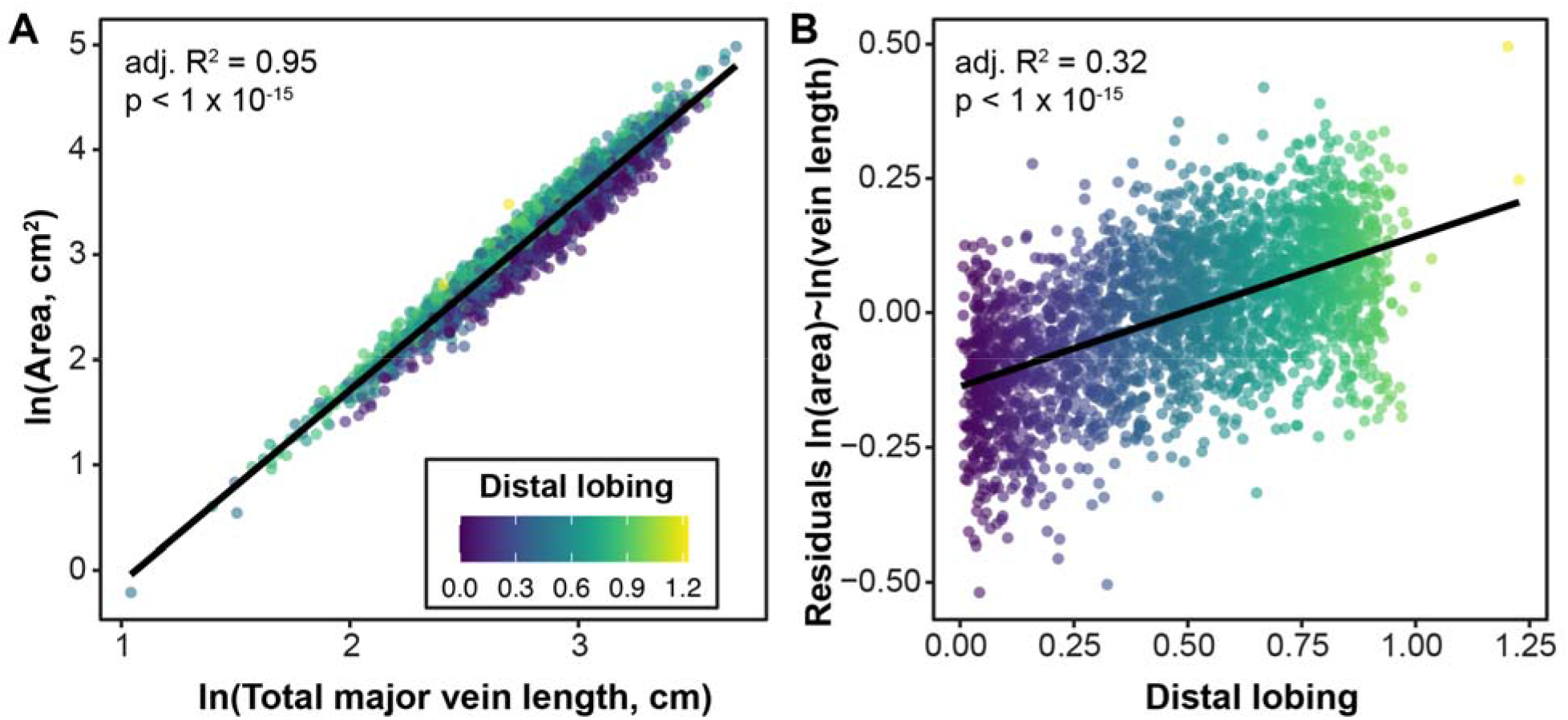
(A) Natural logarithm of area (cm^2^) vs natural logarithm of total major vein length. Total major vein length (cm) calculated by combining the length of the proximal, distal, and midvein (B) Residuals from the model of natural logarithm of area (cm^2^) vs natural logarithm of total major vein length, as indicated in panel A, vs distal lobing. In both panels, each leaf (n=2632) is plotted with the color of the point indicating the distal lobing value. The natural log of leaf area varies significantly as a function of the natural log of total major vein length, and residuals from that model vary significantly as a function of distal lobing. The adjusted R^2^ for each model is indicated on the plots.

In addition to total major vein length, we also calculated total branching vein length across other veins measured in this study, and found that the two measurements were highly correlated (Figure S3, r=0.97, p <1 ×10^−15^). Leaves measured for this study with higher major vein lengths (longer leaves) also had wider lobes, as evaluated using the length of branching veins. A similar linear relationship between the natural log of leaf area and the natural log of branching vein length occurred (Figure S4A, R^2^=0.96, p <1 ×10^−15^) indicating that like major vein length, leaf area increases as a function of branching vein length. However, when we model the residuals from this relationship as a function of distal lobing, the amount of variation in distal lobing explained by these residuals was lower (Figure S4B, R^2^=0.14, p <1 ×10^−15^). Therefore, distal lobing is a better predictor of the residuals for the relationship between leaf area and major vein length, than branching vein length. For leaves with the same surface area, those which are more highly lobed will tend to have both longer major veins and branching veins, but this difference is more pronounced for the major veins, indicating that leaves compensate for area lost to lobing primarily through length of the lobes, and not width of the lobes.

While increasing the length of the leaf is one way for a more lobed leaf to achieve a similar surface area to a more entire leaf, an alternative or complementary hypothesis is that the leaf has a higher vein to blade ratio. Although we did not observe a strong difference in vein to blade across years despite leaves increasing in size (Figure S1), previous work has identified vein to blade ratio as a strong indicator of allometric variation, with larger leaves decreasing the proportion of the area composed of vein relative to blade (Chitwood et al., 2021). By modeling the relationship between the natural log of leaf area and the natural log of vein to blade ratio, we identified a significant linear relationship, with total leaf area decreasing as vein to blade decreased (Figure 4A, R^2^=0.24, p <1 ×10^−15^). This linear, allometric relationship is similar, albeit weaker, than the one seen between area and total major vein length (Figure 3A) and total branching vein length (Figure S4A). However, it occurs in the opposite direction, with leaves having smaller total areas as the vein to blade ratio increases and the total major and branching vein lengths decrease.

**Figure 4.**
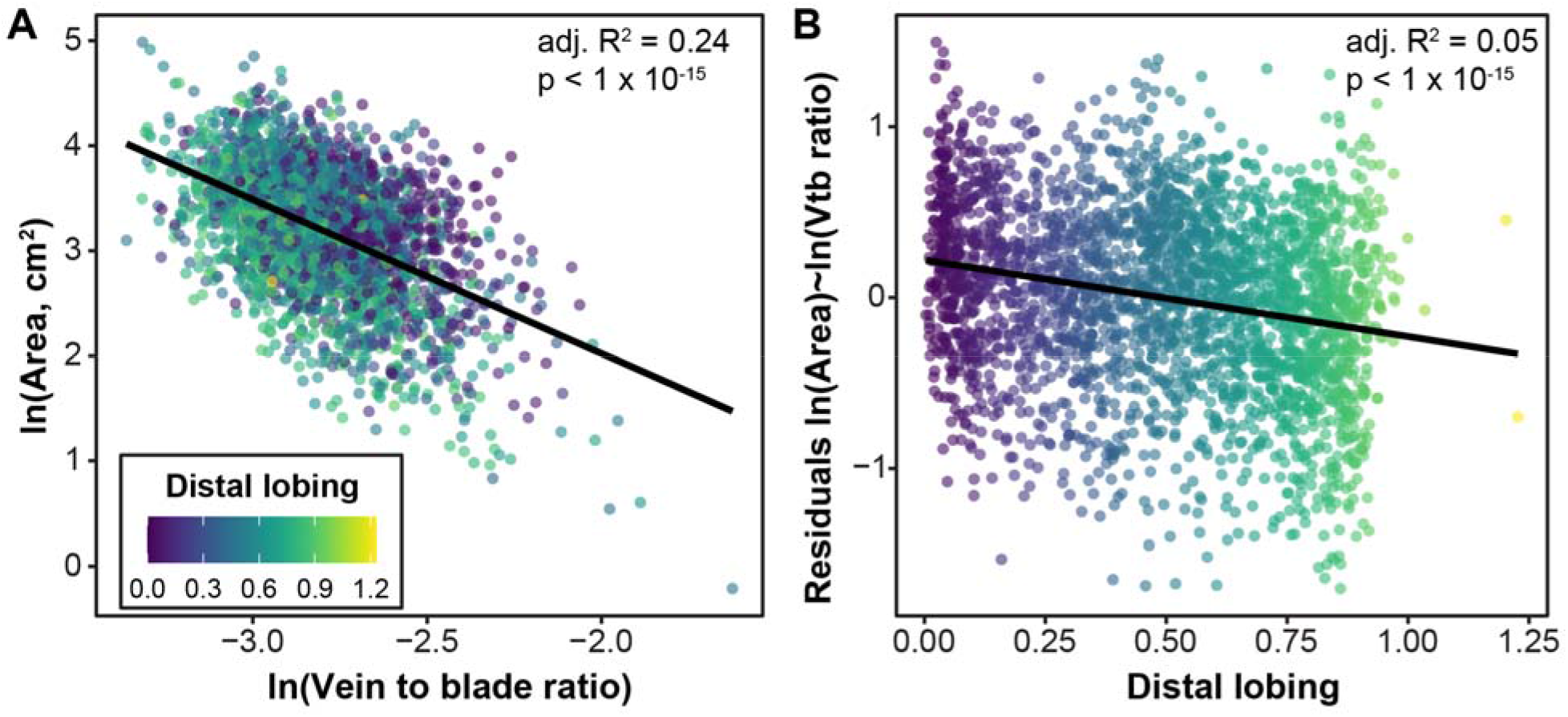
(A) Natural logarithm of area (cm^2^) vs natural logarithm of vein to blade ratio. (B) Residuals from the model of natural logarithm of area (cm^2^) vs natural logarithm of vein to blade ratio, as indicated in panel A, vs distal lobing. In both panels, each leaf (n=2632) is plotted with the color of the point indicating the distal lobing value. The natural log of leaf area varies significantly as a function of the natural log of vein to blade ratio, and residuals from that model vary significantly as a function of distal lobing. The adjusted R^2^ for each model is indicated on the plots.

The residuals from the model between the natural log of leaf area and the natural log of vein to blade ratio explain a significant amount of the variance in distal lobing (Figure 4B, R^2^=0.05, p <1 ×10^−15^), but the R^2^ is much smaller than for the same models applied using total major or branching vein lengths. Thus, leaf area is maintained by more highly lobed leaves primarily through increases in length, not increases in the surface area of veins relative to the blade. However, this subtle relationship still indicates that for leaves of the same size, those that are more highly lobed will have higher vein to blade ratios, which partly explains how they are able to compensate for the reduction in blade area.

## Discussion

By quantifying variation in shape for 2,632 leaves sampled across 476 grapevines, showing immense variation in leaf lobing, we were able to evaluate the relationship between leaf size and leaf shape. Previous work performing linkage mapping in grapevine identified quantitative trait loci on chromosome 1 associated with the depth of the leaf sinus, or “lobiness” (Welter et al., 2007; Demmings et al., 2019). Although we did not perform linkage mapping with the biparental crosses examined in this work, we did identify distal lobing as the primary source of variation. This finding is supported both by the historical literature (Galet, 1979) as well as a detailed study of 60 wine and table grape varieties, which found that the distal sinus was one of the strongest indicators of variety (Chitwood, 2021).

Although lobing was the primary source of variation in shape across the leaves measured, leaf area varied only slightly as a function of lobing, with an R^2^ of 0.003. Indeed, due to the presence of a handful of smaller, more entire leaves, the slope of the linear relationship between the natural log of leaf area and distal lobing was negative (−0.038) indicating that more entire leaves with distal lobing values closer to 1 actually had slightly smaller leaf areas than larger leaves, although as noted, this relationship is very minor. Without other compensating mechanisms, increasing lobing would reduce leaf area. The lack of any substantial relationship between lobing and leaf size indicates the existence of compensating mechanisms that allow lobed leaves to maintain overall area.

We determined that leaf area increases as a function of total major vein length and total branching vein length and leaf area decreases as a function of vein to blade ratio. These relationships are stronger for more highly lobed leaves, with the residuals for each model differing as a function of distal lobing. More highly lobed leaves (with lower distal lobing values) have more negative residuals for total major vein length and total branching vein length, and more positive residuals for vein to blade ratio. These relationships indicate that for a given leaf area, more highly lobed leaves have longer veins and higher vein to blade ratios, allowing them to maintain similar leaf areas despite increased lobing. These findings show how more highly lobed leaves may compensate for what would otherwise be a reduced leaf area, allowing for increased photosynthetic capacity through similar leaf size.

Our analyses here are restricted to five biparental populations, all with a shared parent or grandparent responsible for lobing, DVIT 2876. Lobing in DVIT 2876 is derived from the *V. piasezkii,* in which blade dissection can extend completely to the petiolar junction as a 3 to 5-foliate leaf. The populations also contain lobing introduced from *V. vinifera*. It is possible that the relationships observed here are not necessarily true across *Vitis* more broadly, especially in species with more entire leaves.

Determining how lobing influences variation in other measurements of leaf shape is not just botanically fascinating, but also important because more highly lobed leaves may have a viticultural benefit in the vineyard. For example, in cotton, leaf shape has an impact on chemical spray penetration, with the highly lobed leaves allowing more spray to be delivered deeper into the canopy (Andres et al., 2016). However, in cotton, the switch from entire to highly lobed leaves reduces the leaf area (Andres et al., 2016). In grapevine, extensive canopy management practices such as planting distance, pruning including shoot length and architecture, training system, and trellis design all are used to influence access to sunlight (Reynolds and Vanden Heuvel, 2009; Keller, 2020b). Given the central position of grape berries in the canopy, access to sunlight is particularly important, and leaf removal or thinning may be used to increase sunlight penetration around the canopy fruit zone. Leaves within the canopy may receive only 1/100, or less, of photosynthetically active radiation that exterior leaves do (Smart et al., 1982). Light in the canopy has an important influence on grape composition. For example, in one study of ‘Merlot’ grapes, leaf removal significantly reduced titratable acidity and increased the concentrations of phenols, anthocyanins, flavonols, and flavan-3-ols (Anić et al., 2021). Similarly, ‘Riesling’ grapes had higher monoterpene alcohols and bound aromatic alcohols (Zoecklein et al., 1998) and ‘Sauvignon blanc’ grapes showed increased total soluble solids and reduced titratable acidity (Bledsoe et al., 1988) when leaves were removed. In all of these cases, leaf removal resulted in more light in the canopy fruit zone and the changes in grape composition are attributed to the increase in light.

In addition to grape composition, light is also known to influence fertility of latent buds. Warm, sunny conditions, water and nutrient access, as well as sufficient photosynthesizing leaf area, are all necessary to maximize the number of primordia. Similar conditions are also required for flowering, fruit set, and berry development (Keller, 2020a). Like cotton, access to the grapes within the canopy due to highly lobed leaves or leaf thinning could improve spray penetration and improve the deposition of chemicals. Indeed, there is evidence that grapevine varieties do differ in spray penetration, with previous work examining two grape varieties finding that the vineyard spraying system needed to be calibrated based on variety (Palleja and Landers, 2017).

While leaf thinning may have benefits, it does reduce the leaf area per meter of canopy, by partially or entirely removing leaves. Ultimately, more highly lobed leaves with the same leaf surface area could enable grape growers to capture the benefits of leaf thinning (sunlight and spray penetration) while not removing leaves, thereby not reducing leaf area and associated photosynthetic capacity, as well as saving the management cost of leaf removal.

Across both the paleorecord and extant populations, more highly lobed leaves and smaller leaves have been identified in similar climates (Chitwood and Sinha, 2016). In contrast, within the grapevines studied here, leaf shape and size are decoupled from each other. Rather, increases in vein length compensate for leaf area lost due to lobing, providing evidence of one mechanism which could allow for leaves to maintain area while increasing lobing. This mechanism could allow sunlight to permeate the grapevine canopy, without reducing photosynthetic capacity of the vine.

## Conclusions

By examining the relationship between leaf size and shape, we show that highly dissected leaves do not differ in overall leaf area compared to more entire leaves. Future work is still needed to determine if, whether through traditional breeding or gene editing, such variation in leaf shape could be harnessed for improving management practices of grapevines. These results contrast with previous work performed more broadly across plant families and may suggest use of a novel allometric mechanism in Vitaceae in which reductions in leaf area due to lobing are compensated through increases in vein length.

## Acknowledgments

The research conducted for this study was supported by the National Science Foundation (NSF) Plant Genome Research Program 1546869. JFS was supported by an NSF Graduate Research Fellowship under Grant No. 1758713, and Saint Louis University. This research is also funded through the USDA National Institute of Food and Agriculture and Michigan State University AgBioResearch.

We acknowledge all of the individuals involved in maintaining the vineyard evaluated for this study. We acknowledge Leah Brand (Missouri State University), Julie Curless (Missouri State University), Dalton Gilig (University of Missouri), and Ilona Natsch (Saint Louis University) for assistance in sampling and landmarking of the leaves. We would also like to acknowledge Laszlo Kovacs (Missouri State University) for student supervisory support.

## Data availability

The original scans of leaves used for this study are available from the Dryad Digital Repository: **\*submission in progress\***. All data and code used in this study can be found on github (https://github.com/zoemigicovsky/grape_leaf_lobing).

## Supplemental Figures

**Figure S1.**
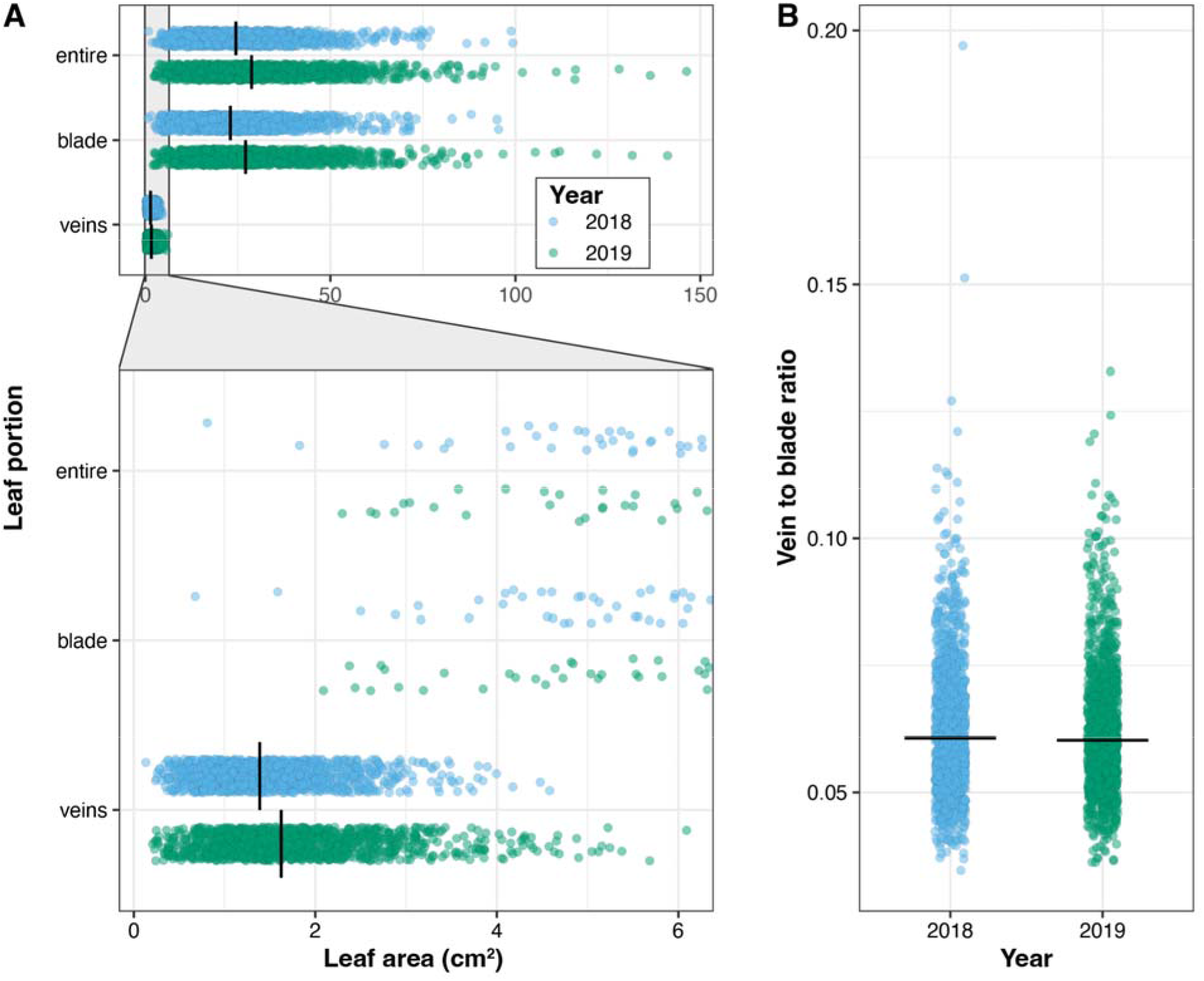
(A) Leaf area (cm^2^) for all leaves sampled in 2018 and 2019. The area values are shown for the whole leaf, as well as the blade and vein portion of the leaves separately. Due to the small values for vein area, a zoomed in plot is shown. (B) The ratio between vein area and blade area for each leaf is shown across 2018 and 2019. Only leaves from vines sampled in both years (vine n=398, leaf n=2388) are shown. The black vertical/horizontal lines indicate the median value.

**Figure S2.**
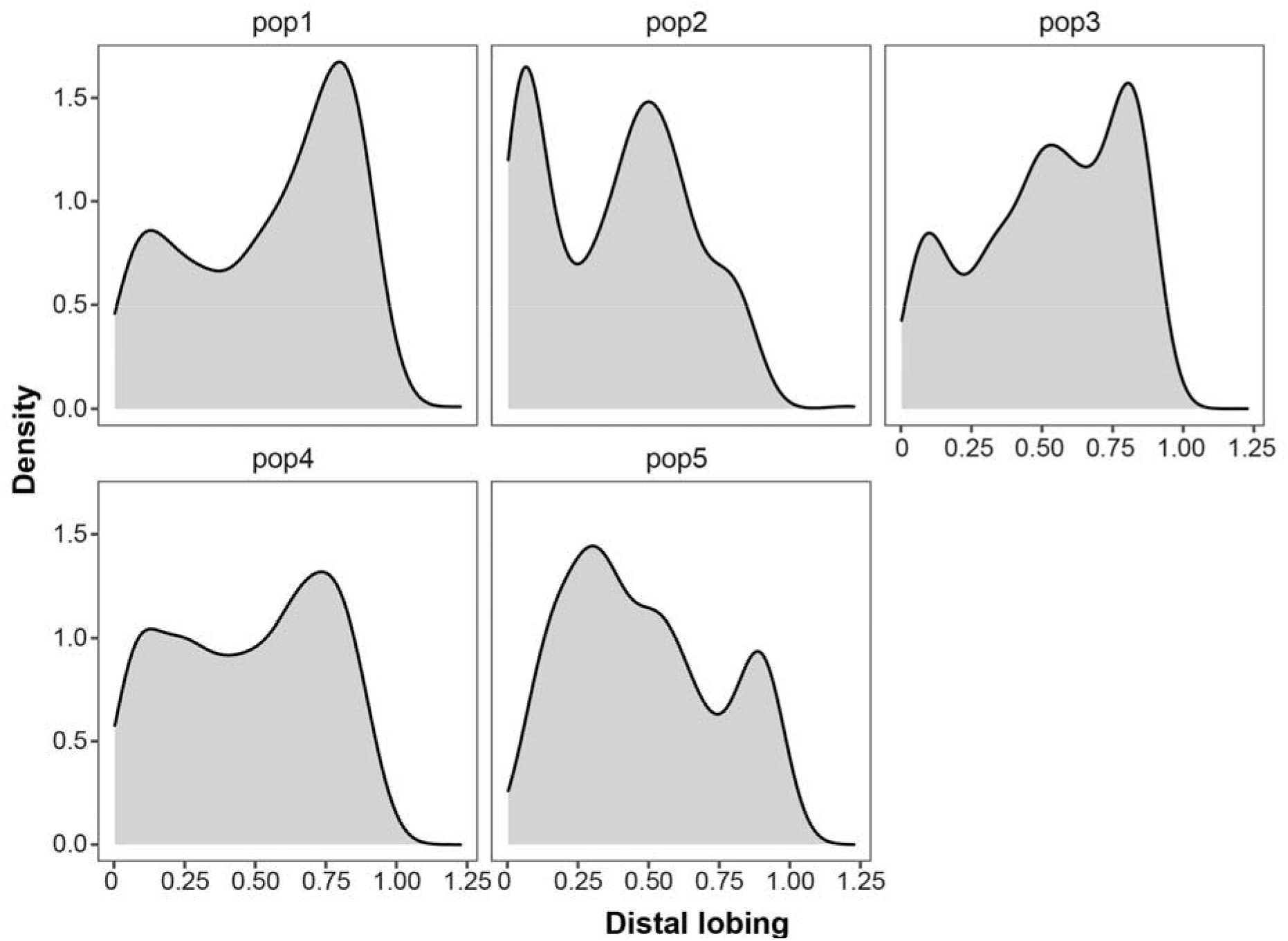
Density plots indicating the distribution of distal lobing values for each of the five populations examined in this study. Distal lobing values are calculated as the ratio of the length of the distal sinus to the length of the distal lobe, with values increasing as lobing decreases.

**Figure S3.**
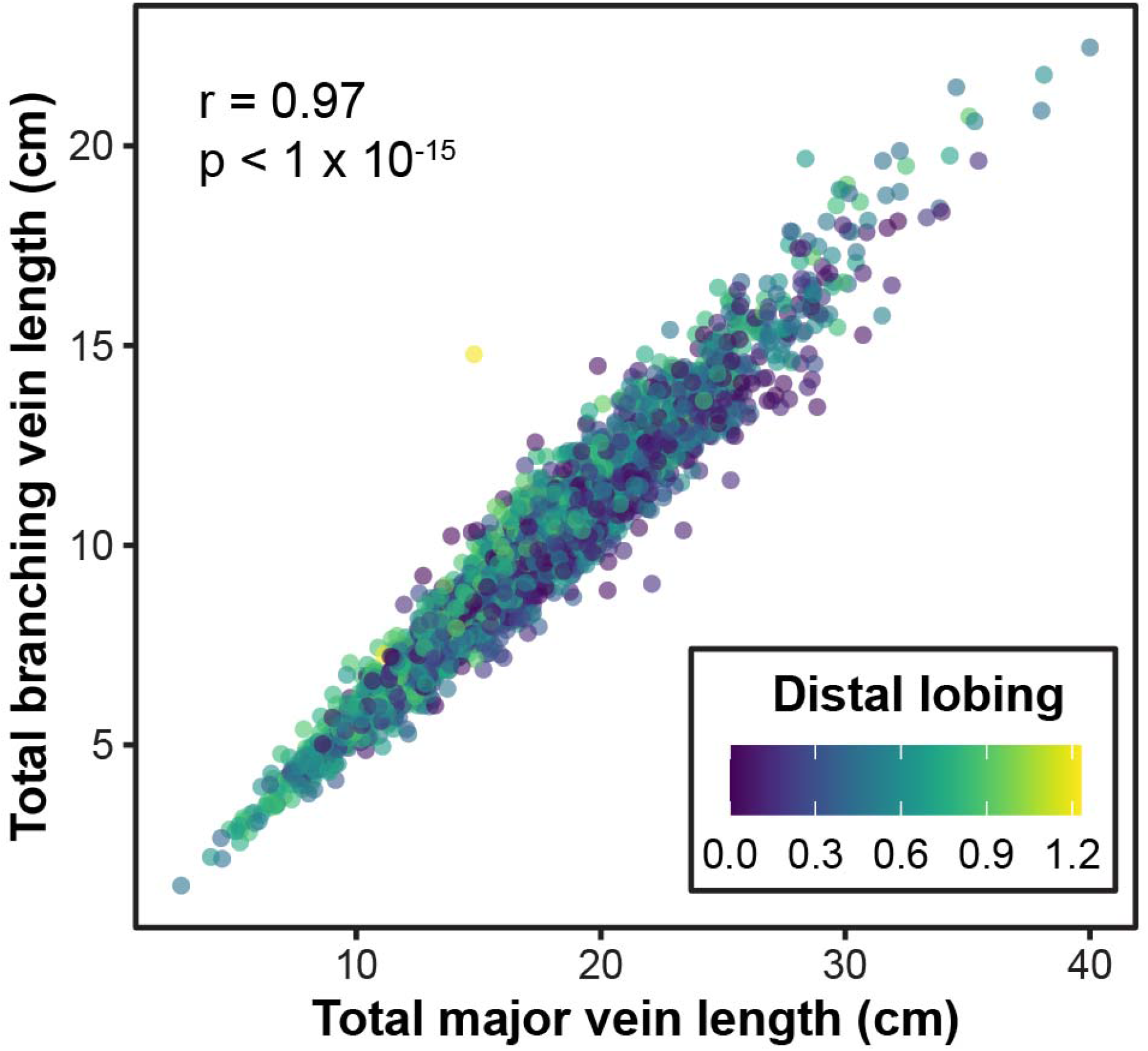
Correlation between the total major vein length (cm) and the total branching vein length (cm). Total major vein length (cm) was calculated by combining the length of the proximal, distal, and midvein. Total branching vein length (cm) was calculated by combining the length of the branching veins shown in Figure 1B, which does not include the proximal, distal, or midvein. Each leaf (n=2632) is shown, with the color of the point indicating the distal lobing value in both panels. The r and p-value of a Pearson’s correlation is indicated.

**Figure S4.**
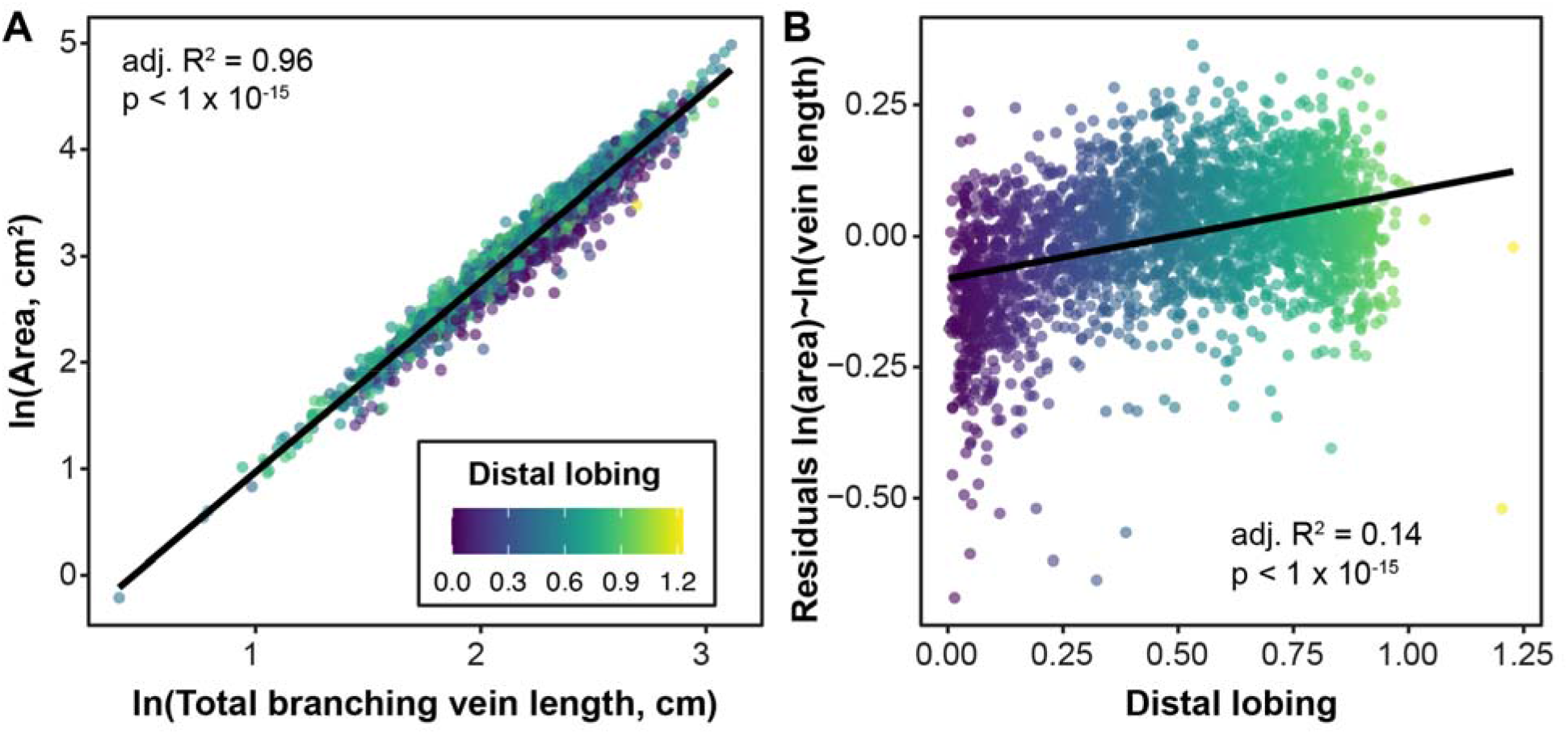
(A) Natural logarithm of area (cm^2^) vs natural logarithm of total branching vein length. Total branching vein length (cm) was calculated by combining the length of the branching veins shown in Figure 1B, which does not include the proximal, distal, or midvein. (B) Distal lobing vs residuals from the model of natural logarithm of area (cm^2^) vs natural logarithm of total branching vein length, as indicated in panel A. In both panels, each leaf (n=2632) is plotted with the color of the point indicating the distal lobing value. The natural log of leaf area varies significantly as a function of the natural log of total branching vein length, and distal lobing varies significantly as a function of the residuals from that model. The adjusted R^2^ for each model is indicated on the plots.

